# T1SEstacker: A tri-layer stacking model effectively predicts bacterial type 1 secreted proteins based on C-terminal non-RTX-motif sequence features

**DOI:** 10.1101/2021.11.10.468166

**Authors:** Zewei Chen, Ziyi Zhao, Xinjie Hui, Junya Zhang, Yixue Hu, Runhong Chen, Xuxia Cai, Yueming Hu, Yejun Wang

## Abstract

The proteins secreted through type 1 secretion systems often play important roles in pathogenicity of various gram-negative bacteria. However, the type 1 secretion mechanism remains unknown. In this research, we observed the sequence features of RTX proteins, a major class of type 1 secreted substrates. We found striking non-RTX-motif amino acid composition patterns at the C-termini, most typically exemplified by the enriched ‘[FLI][VAI]’ at the most C-terminal two positions. Machine-learning models, including deep-learning models, were trained using these sequence-based non-RTX-motif features, and further combined into a tri-layer stacking model, T1SEstacker, which predicted the RTX proteins accurately, with a 5-fold cross-validated sensitivity of ~0.89 at the specificity of ~0.94. Besides substrates with RTX motifs, T1SEstacker can also well distinguish non-RTX-motif type 1 secreted proteins, further suggesting their potential existence of common secretion signals. In summary, we made comprehensive sequence analysis on the type 1 secreted RTX proteins, identified common sequence-based features at the C-termini, and developed a stacking model that can predict type 1 secreted proteins accurately.

## Introduction

Type 1 secretion systems (T1SSs) are uniquely distributed in gram-negative bacteria, which can secrete various substrate proteins through the two bacterial cell membranes by one step (classical) or two steps (non-classical) into extracellular milieu (Spitz et al. 2019; Smith, Sondermann, and O’Toole 2018). A T1SS is composed by three elementary components - an ATP-binding cassette (ABC) transporter located in inner membrane, an outer membrane factor (OMF), and a membrane fusion protein (MFP) connecting the ABC transporter (Kanonenberg et al. 2018). A wide variety of proteins are secreted through this oligomeric secretion channel to play their biological roles. Due to simple structure of the system, T1SSs have been widely applied in biomedical engineering applications (Ryu et al. 2015).

The T1SS substrates, also called Type 1 Secreted Effectors (T1SEs), have various biological function, such as host invasion (virulence factors, e.g. HlyA) (Felmlee, Pellett, and Welch 1985), enzymolysis (digestion enzymes, e.g. TliA and PrtA) (Son et al. 2012), nutrient acquisition (iron-scavenger proteins, e.g. HasA) (Kanonenberg, Schwarz, and Schmitt 2013), biofilm formation (adhesins, e.g. LapA) (Guo et al. 2019), etc. Since the first T1SS substrate, hemolysin A (HlyA), was discovered in 1979 and its nucleotide sequence was determined in 1985 (Noegel et al. 1979; Felmlee, Pellett, and Welch 1985), the structural characteristics and function of T1SEs have been studied extensively. Typical T1SEs can be divided into 3 classes (Classes 1~3) according to different ABC transporters types: C39-containing ABC transporters with hydrolase activity, C39-like domain (CLD)-containing ABC transporters without hydrolase activity, and a third type of ABC transporters without any additional N-terminal domain (Hui et al., 2021). Class 1 T1SEs, known as the smallest T1SS substrates, normally contain N-terminal leader peptides. The C-termini of the leader peptides contain a canonical double glycine (‘GG’) motif, which can be recognized and cleaved by the C39 domains of corresponding ABC transporters before the mature proteins are secreted through T1SSs (Kanonenberg, Schwarz, and Schmitt 2013). Class 2 T1SEs have remarkable repeats-in-toxin (RTX) domains and are also known as RTX proteins. The glycine-rich nanopeptide repeats in RTX domains show a ‘GGxGxDxUx’ consensus sequence motif where ‘x’ is any amino acid and ‘U’ represents a large or hydrophobic amino acid. Class 3 T1SEs may also contain RTX repeat sequences but not necessarily. The last two categories do not contain N-terminal leader peptides, but instead potentially have secretion signal sequences in the C-termini. However, the C-terminal signal patterns and function mechanisms remain to be clarified (Kanonenberg et al., 2013). Recently, a group of non-classical T1SEs named RTX adhesins (Class 4) have been reported, which are closely related to biofilm formation (Smith et al., 2018). Different from Classes 1-3 T1SEs, the RTX adhesins are transported from cytoplasm to extracellular environment by a two-step secretion mechanism, which involves periplasmic intermediates. This subgroup of T1SS machinery is linked with a bacterial transglutaminase-like cysteine proteinase (BTLCP) (Smith et al., 2018). The RTX adhesion proteins have dialanine BTLCP cleavage sites in the N-terminal retention module that can be recognized and cleaved by the machinery-coupled BTLCP in periplasm before the cross-outer-membrane transport (Boyd et al., 2014; Smith et al., 2018). The currently known RTX adhesins also have RTX repeats and signal sequences in the C-termini (Boyd et al., 2014; Smith et al., 2018).

Both the function and sequences of T1SEs show large diversity, and till now only ~100 homology-not-filtered T1SEs have been validated (http://61.160.194.165/TxSEdb). Bioinformatic strategies have also been tried to predict novel T1SEs, but mainly focused on the RTX proteins with the consensus RTX motifs (Linhartova et al., 2010; Luo et al., 2015). For instance, Linhartova et al combined pattern searching, HMM profiles and RPS-BLAST, to predict 1,024 candidate RTX proteins from 840 bacterial genomes, as comprised the most comprehensive list of RTX T1SE candidates (Linhartova et al., 2010). Luo et al made the first attempt to develop a machine-learning model to predict RTX proteins (Luo et al., 2015). The random forest based model learned amino acid sequence-derived features extracted from the full-length and C-terminal sequences of T1SE candidates predicted by Linhartova et al (Luo et al., 2015). Regretfully, neither a software tool nor a webserver was provided for users to implement the method. Besides, both the homology-based and machine-learning methods completely studied on the RTX proteins and the conserved RTX motif was placed with a large weight. The methods are hardly generalized to find more novel T1SEs without RTX motif features.

By sequence pattern analysis carefully, previously, we identified the position-specific amino acid composition (Aac), secondary structure element (Sse) and solvent accessibility (Acc) features of type 3 secreted effectors (T3SEs) within their N-termini and the various Aac, Sse and Acc profiles of type 4 secreted effectors (T4SEs) within their C-termini (Wang et al., 2011; Wang et al., 2014). Given the unsound evidence about the potential C-terminal secretion signals of T1SEs, in this research, we comprehensively observed the amino acid sequence patterns, especially non-RTX-motif features within the C-termini of RTX proteins, and also the Sse and Acc property. Furthermore, we developed machine-learning models to learn the newly observed sequence-derived features and predicted T1SEs with or without typical RTX motifs. Deep learning models and ensemblers have been recently widely used to predict bacterial secretion signals and achieved good performance (Almagro Armenteros et al., 2019; Xue et al., 2019; Wang et al., 2018; Wang et al., 2019; Hui et al., 2020). We also tested Deep Neural Network models and integrated them and others within a stacked model to improve the prediction performance.

## Materials and Methods

### 1. Datasets

Bacterial RTX proteins were collected from Linhartova et al., 2010. In total, there were 1,024 RTX proteins predicted from 840 bacterial genomes (Linhartova et al., 2010). CD-HIT was used to detect homology among the RTX proteins, while 30% was considered as the similarity cutoff and only one representative was retained if there were multiple proteins showing sequence similarity above the cutoff (Li and Godzik 2006). Proteins were also sampled randomly from the whole proteomes derived from various bacterial genome sequences. The known T1SEs, RTX proteins and their homologs with >30% blastp similarity were removed, and a homology filtering strategy similar to that applied for RTX proteins were used to identify the non-redundant non-RTX proteins. In total, 512 non-redundant RTX proteins were remained, which were considered as the positive dataset (p). 2000 proteins were also randomly selected from the processed non-RTX proteins, and three groups, each with 512 proteins, were further picked out to match the number and general length distribution of the RTX proteins, forming the negative datasets (n1~n3). The p and n1 were used as the main observation datasets. A 5-fold cross-validation strategy was used for training the machine-learning prediction models, for which both the positive and negative datasets were split into 5 subsets of equal size of protein sequences, with four of them being served as training datasets and the rest one as testing datasets in each fold of model analysis. Experimentally validated T1SEs were also annotated manually from literature. These proteins could be RTX or other type of proteins with experimental evidence to be transported through T1SSs. All the datasets were together with the standalone T1SEstacker package and publically available (see ‘Software Availability’; http://61.160.194.165/TxSEdb).

Once the datasets were collected and annotated, the sequence-based features were analyzed with in-house scripts. The secondary structure and solvent accessibility were predicted with SSpro/ACCpro5, with 3 elements encoded for secondary structure (‘H’ for helix, ‘E’ for strand, and ‘C’ for coil) and 2 elements for accessibility (‘B’ for ‘buried’ and ‘E’ for ‘exposed’) (Magnan and Baldi 2014).

### 2. Sequential and Position-specific Amino Acid Composition Features Based Non-Deep-Learning Models

The number and position distribution of RTX motifs featured as ‘GGxGxD’, was observed within the RTX and non-RTX proteins. Sequential Aac, continuous and 1 or 2 amino acid interrupted bAac features, were extracted from the C-terminal 20 or 60 -residue fragments of positive and negative datasets, observed and compared. The features were used for training Random Forest (RF), Support Vector Machine (SVM) and Naive Bayesian (NB) models, with R packages of ‘randomForest’, ‘e1071’ and the ‘e1071’ method ‘naiveBayes’ respectively (https://www.r-project.org/). The neighbor-position Aac conditional constraint features in the C-termini were learned in Markov models (Wang et al., 2013). Bi-Profile Bayesian Position-specific Aac features were extracted, and trained with SVM models (Wang et al., 2011). For the SVM models, four kernels (‘linear’, ‘polynomial’, ‘sigmoid’ and ‘radial’) were tested and the corresponding parameters, e.g., *gamma* and/or *cost*, were optimized using a 10-fold cross-validation grid search strategy within each training dataset. For the other models, the features were also extracted based on each training dataset. The details about the models and the optimized parameters refer to the website of T1SEstacker (see ‘Software Availability’).

### 3. Deep Learning Models

Deep learning models were trained with the Aac features of RTX proteins within the C-terminal 20 (C20) and 60 amino acid positions (C60). Each position was represented by a 20-element feature vector describing the composition of amino acids. A *m* x 20*L* matrix was built to represent the original Aac features of training datasets, where *m* was the number of training proteins and *L* was 20 or 60 for C20 or C60 models respectively. Fully connected Deep Neural Network (DNN), Self Attention (SelfAttention) and models with Long-Short Term Memory (LSTM) cells (RNN) were trained and tested with a 5-fold cross-validation strategy. The details about the models and the optimized parameters refer to the website of T1SEstacker (see ‘Software Availability’).

### 4. A Stacked Model Featured By The Prediction Results of Individual Models

To achieve better prediction performance, we proposed a new stacking scheme to integrate prediction results of individual models (Fig. 1). A primary stacked model was built for each original 5-fold training dataset and its based individual models. For each original 5-fold testing dataset, an embedded 5-fold cross validation was adopted to evaluate the performance of stacked models. The prediction result of each-fold best-trained model of individual algorithms on each protein of the corresponding testing dataset was based, and encoded as 1 (RTX) or 0 (non-RTX) according to the model-specific optimized cutoff score. Each protein within an embedded 5-fold training dataset was represented as a feature vector of ‘0’ and ‘1’, and an *m’* x *n* matrix was generated for the whole training dataset, where *m’* is the protein number of the embedded training dataset and *n* is the number of individual machine-learning models. SVM models with ‘linear’ kernels were trained and the parameters (costs) were optimized with a 10-fold cross-validation grid-searching strategy.

**Fig 1.**
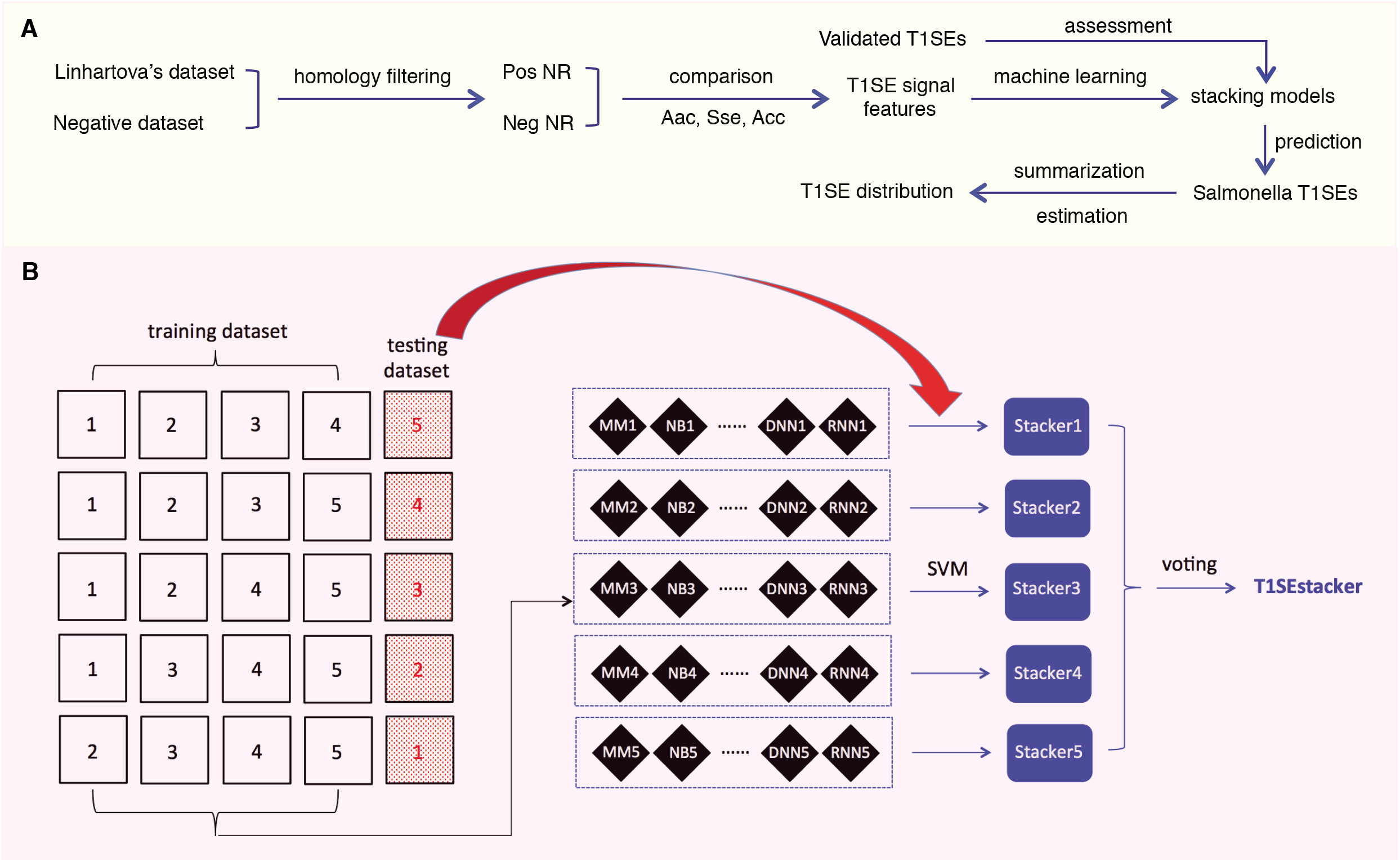
Research design and scheme of the tri-layer staking model T1SEstacker. **(A)** Research design. Pos, positive; Neg, negative; NR, non-redundant. **(B)** Tri-layer stacking model. The original positive and negative datasets were five-fold divided and four of them were used as training datasets for training different machine-learning models, e.g., MM, NB, etc. The remained sub-divided dataset was used as the corresponding testing dataset. For each round of cross validation, the prediction results of the individual models for the testing dataset were used for training and testing the primary stacker models using SVM with an embedded 5-fold cross-validation strategy. Prediction results of the primary stackers were further stacked in T1SEstacker with a voting strategy.

A voting strategy was used to integrate the five primary stacked models, with the same weight assigned for each model.

### 5. Performance Evaluation of The Individual and Stacked Models

Sensitivity (Sn), Specificity (Sp), Accuracy (ACC), the Area Under the Curve of Receiver Operating Characteristic (rocAUC) and Matthews Correlation Coefficient (MCC) were defined as below and used as measures to assess the performance of models based on a 5-fold cross-validation strategy.

Sn = TP / (TP + FN)
Sp = TN / (TN + FP)
ACC = (TP + TN) / (TP + FN + TN + FP)
MCC = ((TP x TN) – (FN x FP))/sqrt((TP+FN)x(TN+FP)x(TP+FP)x(FN+FN))

TP, TN, FP and FN denote the number of true positives, true negatives, false positives and false negatives, respectively.

### 6. Statistics

Individual amino acids were counted within C-terminal 20, 60 or 110-aa fragments, and Mann-Whitney tests were performed to compare their distribution between RTX and non-RTX proteins, followed by Bonferroni corrections. For continuous or non-continuous bi-AAs, the composition was also compared between the C-termini of RTX and non-RTX proteins using the same statistical methods. Another balanced rate comparison method, EBT, was also adopted to compare the C-terminal occurrence of bi-AAs between the two classes of proteins (Hui et al., 2017). The alpha levels for all tests were preset as 0.05.

### 7. Software Availability

T1SEstackers and its modules were developed with Python, Perl and R. The packages and user manual can be downloaded freely via the link, http://www.szu-bioinf.org/tools/T1SEstacker. A web server was also initiated to make internet-based prediction service: http://www.szu-bioinf.org/T1SEstacker.

### 8. *Salmonella* genomes

In total, 26 representative strains were included, which covered the known *Salmonella* phylogenetic groups. N268_08, NCTC12419 and RKS3044 belong to S. *bongori*, RKS2983 and RSK2980 belong to *S. enterica subsp*. *arizonae*, ATCC_BAA_1581 and RKS3027 belong to *S. enterica subsp*. *houtenae*, 2439-64 and RKS3013 belong to *S. enterica subsp. vii*, 11_01853, 11_01854, 11_01855 and RKS2978 belong to *S. enterica subsp*. *diarizonae*, RKS2986 and ST114 belong to *S. enterica subsp*. *salamae*, 1121 and RKS3057 belong to *S. enterica subsp*. *indica*, while P12519, 287/91, ATCC9150, SPB7, RKS4594, ATCC9120, CT18, 14028S and LT2 represent various serovars of *S. enterica subsp*. *enterica*. The genome and genome-encoding proteome were downloaded from NCBI genome database (https://www.ncbi.nlm.nih.gov/genome). T1SEstacker was applied to predict the T1SE candidates with default settings.

## Results

### 1. Research design

The major obstacles for training machine-learning models in prediction of bacterial T1SEs include (1) the limited number of experimentally validated positive proteins, and (2) the large sequence diversity of T1SE groups. Comprehensive literature searching and manual annotation only curated 99 validated T1SEs, and only 49 were retained after a strict homology-filtering process, which were distributed in all the four major T1SE groups (http://61.160.194.165/TxSEdb). To better analyze the likely novel sequential features that could facilitate understanding the mechanisms of type 1 secretion and prediction of new T1SEs, and as performed by others previously (Luo et al., 2015), we took the larger-scale RTX T1SE candidates identified by Linhartova et al., 2010 as training data for analysis of features other than RTX motifs and building models to predict novel T1SEs.

After removing the homologs, the remained non-redundant T1SEs and paired non-T1SEs were compared for their sequential and position-specific Aac, Sse and Acc features, especially non-RTX motif features (Fig. 1A). With the sequence-based features, a stacking model was developed to predict T1SEs (Fig. 1B). Representative strains of *Salmonella* phylogenetic branches were predicted with the newly developed model, and the possible number and distribution of candidate T1SEs were evaluated (Fig. 1A).

### 2. Distance distribution of RTX motifs to the C-termini in RTX proteins

The 512 non-redundant RTX proteins show a length distribution from 70 to 36,805 amino acids, with a median of 1,112 residues and 7 super-long proteins with larger than 10,000 amino acids (Fig. 2A). 494 from the 512 positive proteins could be found with at least one RTX motif within each protein sequence (Fig. 2B). As a control, only 13 from the total 2,341 non-redundant negative proteins contained RTX motifs, which were filtered for further comparative or model-training analysis. The most C-terminal residue of each most C-terminal RTX motif shows a distance of 1 – 21,948 amino acids to the C-terminus of the corresponding full-length protein, with a median of 110 amino acids (Fig. 2C). Fewer than 9 percentages of the C-terminal RTX motifs have a distance of smaller than 60 amino acids from the protein C-termini, and only ~5 percentages are shorter than 20 amino acids (Fig. 2D).

**Fig 2.**
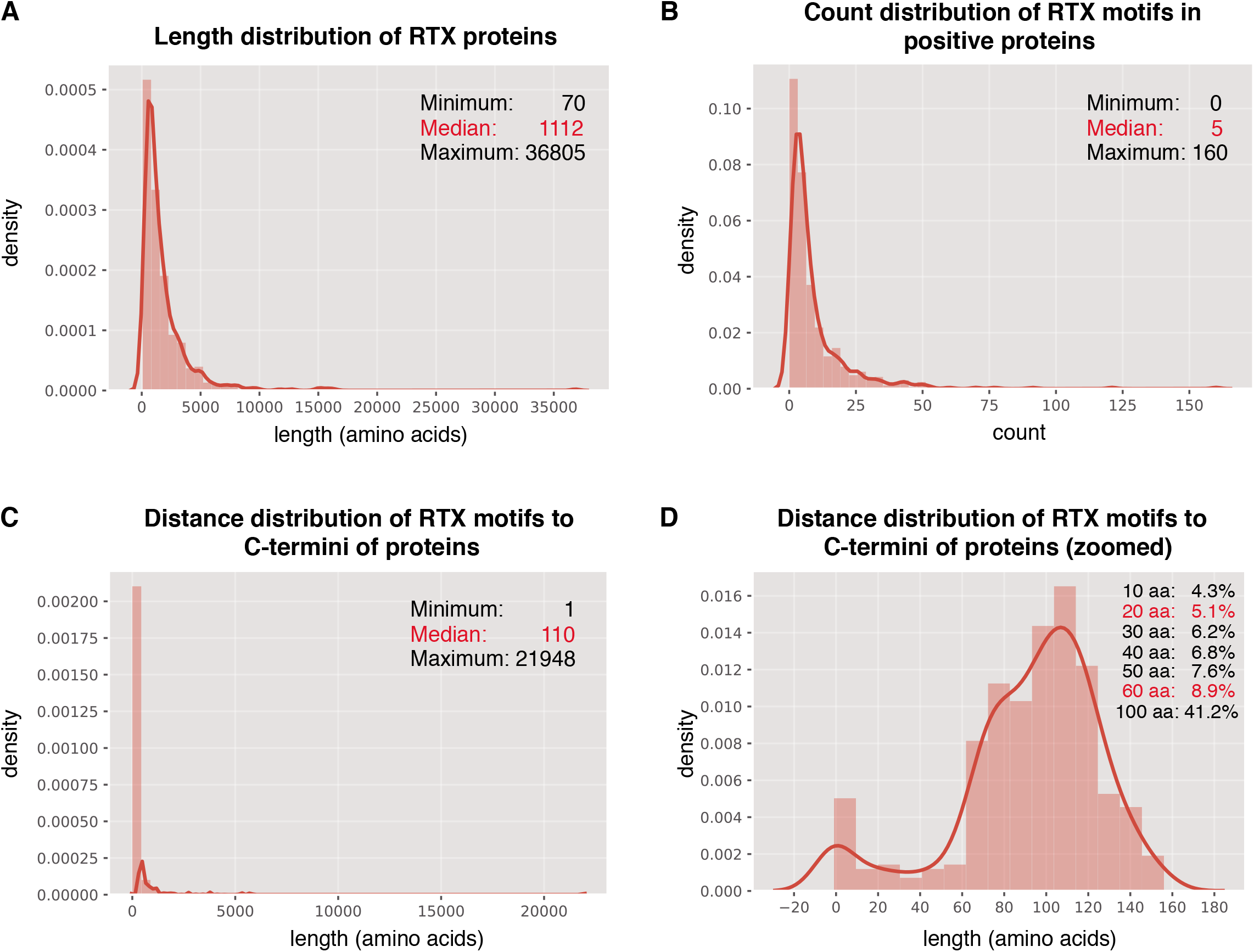
Distribution of RTX motifs in RTX proteins. **(A)** Length distribution of the RTX proteins. **(B)** Count distribution of RTX motifs in the RTX proteins. **(C)** – **(D)** Distance distribution of RTX motifs to the C-terminal ends of the RTX proteins. The accumulated probabilities of the proteins with the RTX motif distance of ≤10~60-aa and 100-aa from the C-termini were shown in **(D).**

### 3. Sequential amino acid composition features buried in the C-termini of RTX proteins

We compared the composition of individual amino acids (Aac) and two continuous or non-continuous amino acids (bAac) among the C-termini of RTX proteins, since there were possibly atypical secretion signals (Boyd et al. 2014; Smith et al. 2018). To avoid the possible misinterpretation caused by RTX motifs, we mainly observed the Aac and bAac profiles within the C-terminal 20 (C20) and 60 residues (C60) (Supplemental Dataset S1). Within C20, most individual amino acids show different composition between the positive and negative proteins, with Aspartic acid (D), Leucine (L), Threonine (T), Valine (V), Isoleucine (I) and Phenylalanine (F) being most typically enriched and Arginine (R), Lysine (K), Glutamic acid (E) and Proline (P) being most strikingly depleted in RTX proteins (Fig. 3A; Mann-Whitney *U* tests with Bonferroni correction, *p* < 0.001). Glycine (G) was not different between the two types of proteins (Fig. 3A; *p* = 1). When the observed length increases to C-terminal 60-aa, most of the featured residues identified from shorter fragments remain different between groups for the composition, whereas some others start to show difference or no difference, e.g., ‘G’ being enriched in RTX proteins and ‘L’, ‘V’ and ‘I’ becoming no difference (Fig. 3A). The enrichment of ‘G’ in RTX C60 fragments is not likely due to the increasing occurrence of RTX motifs which is enriched with ‘G’, since the RTX motifs are lowly represented and the RTX motif featured ‘GG’ is either not strikingly higher in the C60 fragments of RTX proteins (Fig. 2D; Fig. 3B). For the C-terminal 110-aa fragments, the amino acid species with significantly different composition and the amplitude of difference further increase (Fig. 3A). It cannot be excluded that the increased number of RTX motifs leads to the most striking composition amplitude change of ‘D’ and ‘G’. However, the ‘L’ composition change is interesting, which shows higher composition in C20, no difference in C60 and lower composition in C110 of RTX proteins (Fig. 3A).

**Fig 3.**
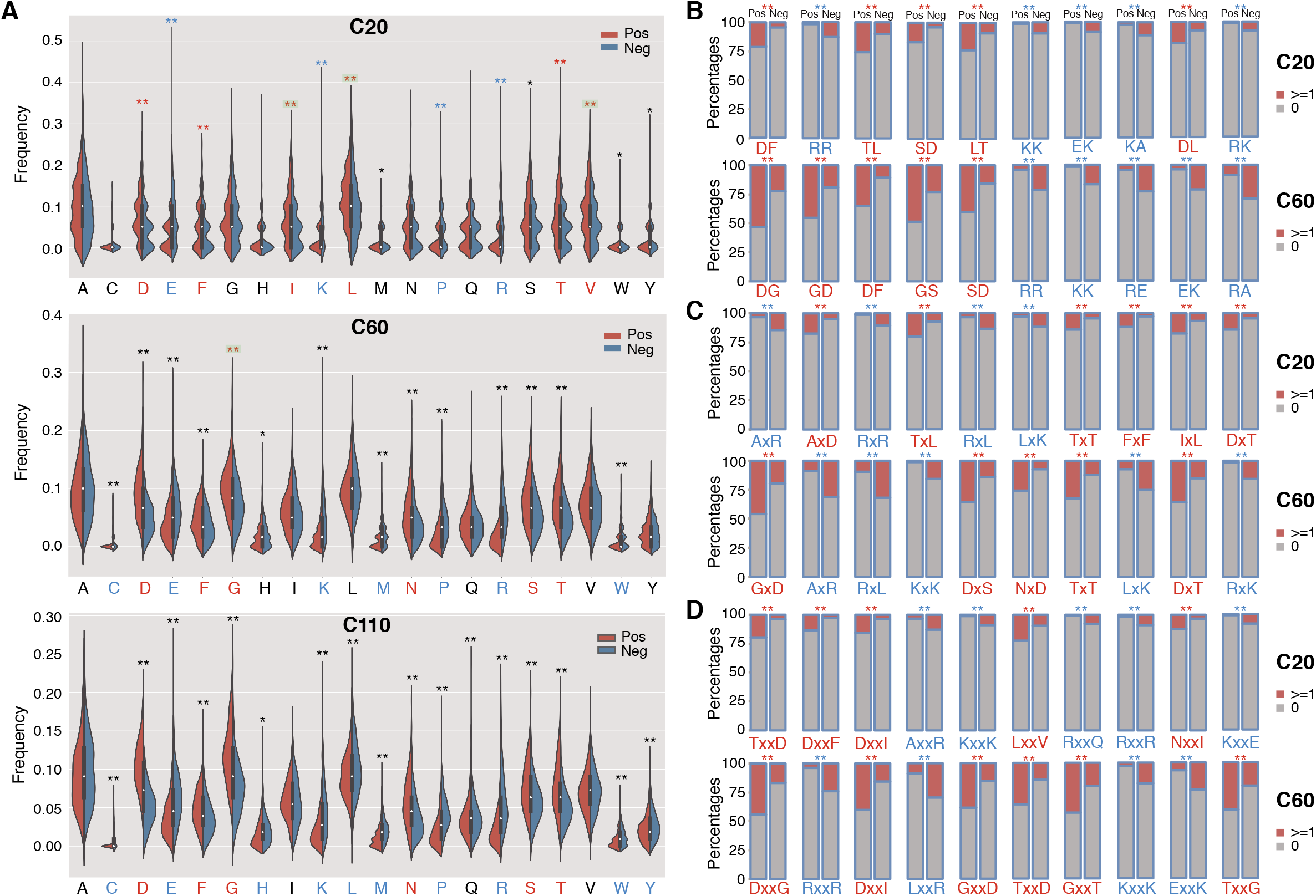
The composition profile difference between the C-termini of RTX and non-RTX proteins for single AAs, continuous and interrupted bi-AAs. **(A)** Single AAs. Bonferroni-corrected Mann-Whitney tests were performed. **(B)** Continuous bi-AAs. **(C)** One amino acid interrupted bi-AAs. **(D)** Two amino acids interrupted bi-AAs. EBT tests were performed for the bi-AAs. Most significantly enriched or depleted single AAs or bi-AAs in RTX C-termini were shown in red and blue, respectively. The single AAs with biased composition changing among the C-termini of different length were shown with green background. **, p < 0.0001; *,p < 0.05.

The continuous and interrupted bAac profile also shows difference in C-termini between RTX and non-RTX proteins. For example, ‘D[FL]’, ‘TL/LT’, ‘AxD’, ‘Tx[LT]’ ‘TxxD’ and ‘Dxx[FI]’ most frequently occur, whereas ‘R[RK]’, ‘K[KA]’, AxR’, ‘Rx[RL]’, AxxR’, ‘Kxx[KE]’ and ‘Rxx[QR]’ are most strikingly depleted in the C-terminal 20-aa fragments of RTX proteins in contrast to non-RTX proteins (Fig. 3B-D; Mann-Whitney *U* tests with Bonferroni correction, *p* < 0.001; EBT_*p* < 0.001). As the observed C-terminal length increases (to 60 aa), the general bAac profile difference between RTX and non-RTX proteins remains or becomes more typical, with only a few changes. The main changes involve the reduced ‘L’ and increased ‘G’ combinations in the RTX C60 enriched list (Fig. 3B-D). It is noted that, either ‘GG’ or ‘GxG’, which is supposed to be highly represented by RTX motifs, does not show the most significant different composition or occurrence in C60 between RTX and non-RTX proteins, suggesting that the observed different ‘G’-combination composition are not due to the increased percentages of RTX motifs in C60 of RTX proteins. In C110, however, the composition show striking difference for both ‘GG’ and ‘GxG’ between RTX and non-RTX proteins (Supplementary Dataset S1).

Other independent non-RTX proteins datasets are also paired and the profile difference for Aac and bAac in C-termini between RTX and non-RTX proteins shows large consistence.

### 4. Position-specific amino acid composition features buried in the C-termini of RTX proteins

The C-terminal position-specific amino acid composition (psAac) profiles were also compared between RTX and non-RTX proteins. Generally, RTX proteins show much larger amino acid composition preference (Fig. 4A). C20 and C21-60 in RTX proteins also show different preference profiles. C20 shows apparent preference for nonpolar ‘L’ and ‘A’ while C21-60 more prefers polar ‘G’ (Fig. 4A). ‘D’, ‘S’ and ‘T’ are preferred in both C20 and C21-60 of RTX proteins. The results are consistent with and explain the observations on sequential Aac and bAac in C-termini of RTX and non-RTX proteins. Remarkably, the C-terminal endmost two positions in RTX proteins show the most typical psAac bias, with a pattern of nonpolar hydrophobic ‘[FLI][VAI]’ motif (Fig. 4A).

**Fig 4.**
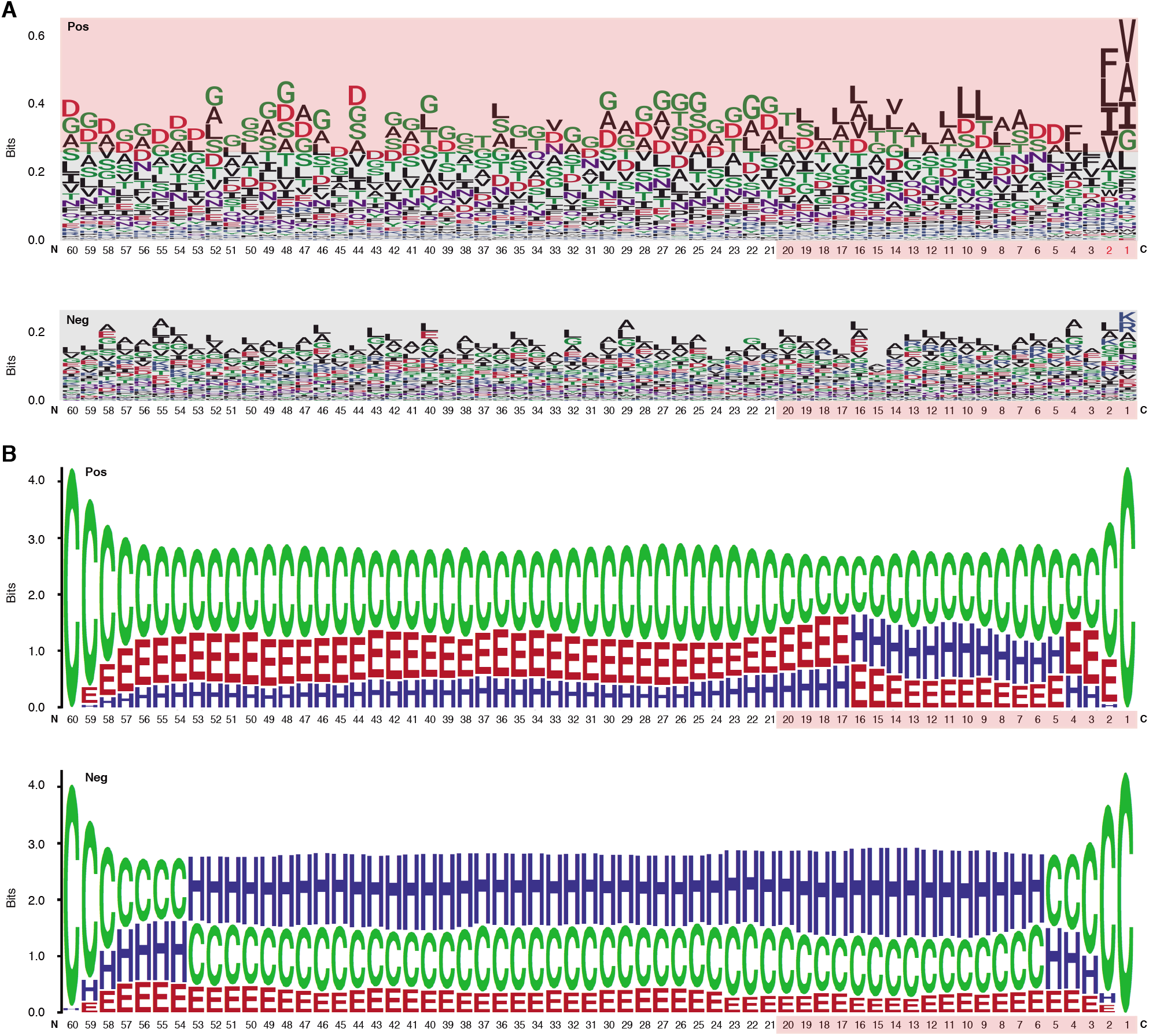
Position-specific Aac and Sse profile difference between the C-termini of RTX and non-RTX proteins. **(A)** Position-specific Aac profile of RTX and non-RTX proteins at C-terminal 60 positions. The strikingly specific bias Aac preference profile of RTX proteins with C-termini and the C-terminal 20 positions of both RTX and non-RTX proteins were shown with pink background. The endmost C-terminal two positions with most typical sequence patterns were shown in red. **(B)** Position-specific Sse profile of RTX and non-RTX proteins at C-terminal 60 positions.

The psAac profile of C-termini of RTX proteins and the difference between them and non-RTX proteins were confirmed with other, paired, independent negative datasets (Supplemental Fig. S1). We also compared the psAac profile of N-termini of RTX and non-RTX proteins (Supplemental Fig. S2). Despite the small difference, it was much more atypical compared to C-termini. Moreover, till now there is no evidence suggesting the existence of type 1 secretion signals within N-termini of the substrate proteins. Therefore, the N-termini were not further studied in this study.

### 5. Enrichment of β-strands and depletion of α-helices within the C-termini of RTX proteins

An apparent difference between the C-termini of RTX T1SEs and non-T1SEs was the depletion of α-helices or enrichment of β-strands and coiled coils, no matter in C20 or C60 (Fig. 4B). The solvent accessibility was not different between the RTX and non-RTX proteins within the C-termini (data not shown). The different forms of secondary structure are likely related with the composition preference of residues. For instance, both polar ‘G’ and nonpolar ‘A’ are enriched in β-strands, while ‘F’ and ‘I’ are not for beneficial for maintenance of the stability of α-helices (Fig. 4A). It remains to be clarified whether the residue composition and structure features are associated with specific recognition of the proteins for specific type 1 secretion.

### 6. C-terminal non-RTX motif features accurately classify RTX from non-RTX proteins

A list of machine-learning models were trained to learn the sequence-based non-RTX motif features buried within the C-termini of RTX proteins, including NB, RF and SVM models learning sequential Aac and bAac features, MM models using adjacent amino acid dependent Aac features, and SVM models analyzing position-specific Aac features (Table 1). Moreover, five types of DL models were trained, with 3 among them of best performance retained (DNN, Attention and RNN), which also learned the C-terminal Aac features of RTX proteins (Table 1). Secondary structure features were not learned in the models since they are not stable, which were predicted with varied accuracy using different software tools.

**Table 1.**
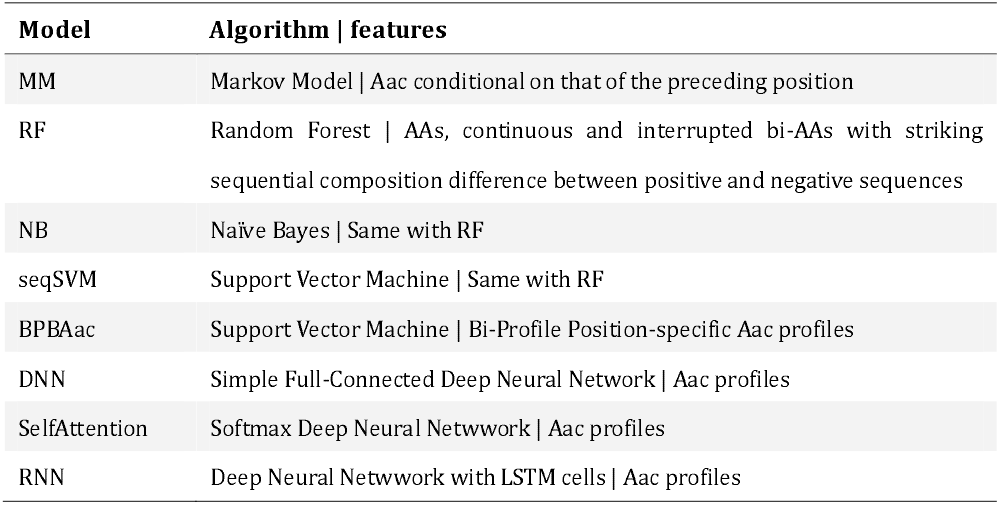
Models and the optimized parameters.

| Model | Algorithm \| features |
| --- | --- |
| MM | Markov Model \| Aac conditional on that of the preceding position |
| RF | Random Forest \| AAs, continuous and interrupted bi-AAs with striking sequential composition difference between positive and negative sequences |
| NB | Naïve Bayes \| Same with RF |
| seqSVM | Support Vector Machine \| Same with RF |
| BPBAac | Support Vector Machine \| Bi-Profile Position-specific Aac profiles |
| DNN | Simple Full-Connected Deep Neural Network \| Aac profiles |
| SelfAttention | Softmax Deep Neural Network \| Aac profiles |
| RNN | Deep Neural Network with LSTM cells \| Aac profiles |

All the models showed certain ability to classify RTX proteins from the non-RTX ones correctly only based on the Aac features within C-terminal 20-aa peptide fragments of known RTX proteins (Table 2; Fig. 5A). RNN, MM, RF, seqSVM showed best prediction performance with the same average rocAUV of 0.88, while BPBAac and DNN appeared poorest with a rocAUV of 0.85 (Table 2; Fig. 4A). C60 models outperformed C20 ones obviously, and MM, RF and seqSVM remained the best-performed models, reached a rocAUC of 0.94 (Table 2; Fig. 5B).

**Fig 5.**
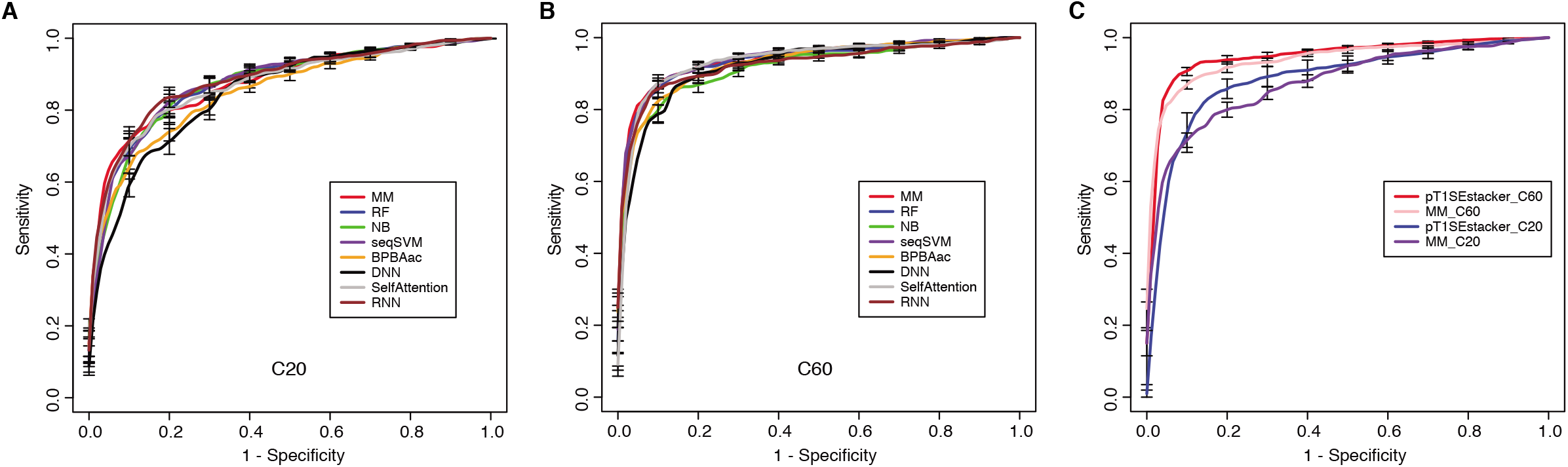
Performance of individual and stacking models on prediction of RTX proteins. **(A)** 5-fold cross-validation ROC curves of individual machine-learning models predicting RTX and non-RTX proteins based on the C-terminal 20-aa features. **(B)** 5-fold cross-validation ROC curves of individual machine-learning models predicting RTX and non-RTX proteins based on the C-terminal 60-aa features. **(C)** Performance comparison of the primary stacking models (pT1SEstacker_C20 and pT1SEstacker_C60) and the representative individual machine-learning models (MM_C20 and MM_C60), based on the average 5-fold cross-validation results.

**Table 2.** Performance of models.

| Model | SN | SP | Acc | rocAUC | MCC |
| --- | --- | --- | --- | --- | --- |
| MM_C20 | 0.81±0.06 | 0.81±0.05 | **0.81±0.02** | **0.88±0.02** | 0.62±0.04 |
| RF_C20 | 0.79±0.06 | **0.82±0.09** | 0.80±0.06 | **0.88±0.04** | 0.61±0.12 |
| NB_C20 | **0.89±0.05** | 0.69±0.06 | 0.79±0.04 | 0.87±0.03 | 0.59±0.09 |
| seqSVM_C20 | 0.79±0.05 | 0.82±0.06 | **0.81±0.04** | **0.88±0.04** | 0.61±0.09 |
| BPBAac_C20 | 0.72±0.06 | **0.82±0.02** | 0.77±0.03 | 0.85±0.03 | 0.55±0.06 |
| DNN_C20 | 0.77±0.05 | 0.75±0.05 | 0.76±0.04 | 0.85±0.03 | 0.53±0.07 |
| SelfAttention_C20 | 0.80±0.03 | 0.80±0.05 | 0.80±0.04 | 0.87±0.02 | 0.60±0.07 |
| RNN_C20 | 0.82±0.05 | 0.80±0.05 | **0.81±0.04** | **0.88±0.04** | **0.63±0.07** |
| pT1SEstacker_C20 | 0.83±0.06 | **0.85±0.04** | **0.84±0.04** | **0.88±0.06** | **0.69±0.09** |
| MM_C60 | 0.86±0.06 | **0.93±0.04** | **0.89±0.02** | **0.94±0.02** | **0.79±0.02** |
| RF_C60 | 0.85±0.06 | 0.90±0.02 | 0.88±0.03 | **0.94±0.03** | 0.76±0.05 |
| NB_C60 | 0.86±0.03 | 0.83±0.05 | 0.84±0.04 | 0.92±0.02 | 0.69±0.07 |
| seqSVM_C60 | 0.84±0.08 | 0.92±0.02 | 0.88±0.04 | **0.94±0.02** | 0.77±0.07 |
| BPBAac_C60 | 0.84±0.04 | 0.89±0.02 | 0.87±0.02 | 0.93±0.01 | 0.73±0.03 |
| DNN_C60 | 0.87±0.06 | 0.85±0.04 | 0.86±0.02 | 0.92±0.02 | 0.72±0.04 |
| SelfAttention_C60 | **0.89±0.02** | 0.89±0.03 | **0.89±0.02** | 0.93±0.01 | 0.78±0.03 |
| RNN_C60 | 0.85±0.07 | 0.90±0.07 | 0.87±0.03 | 0.93±0.03 | 0.75±0.06 |
| pT1SEstacker_C60 | **0.89±0.04** | **0.94±0.02** | **0.91±0.02** | **0.95±0.02** | **0.83±0.03** |

Taking together, the results demonstrate that the C-termini of RTX proteins contain non-RTX Aac signals, which can be used to recognize RTX proteins accurately. The signals are likely distributed along the C-terminal 60-aa positions.

### 7. A stacked model shows striking performance improvement in prediction of RTX types of T1SEs

To achieve better performance, we designed a tri-layer stacking model, which integrates the prediction results of individual models learning sequence-based features, to classify RTX and non-RTX proteins (Fig. 1). The primary SVM-based stacked models (pT1SEstacker) trained with the prediction results of original 5-fold cross-validated testing datasets showed better performance than individual models for both C20 and especially C60, with average rocAUC of 0.85 and 0.95 respectively (Table 2; Fig. 5C). The prediction results of the primary stacked models based on cross-validated testing datasets were assembled in the final model (T1SEstacker) with a voting strategy. It is noted that, with an independent dataset, which will be explained in the next section, we showed that the voting-based tri-layer stacker T1SEstacker generally balanced the effect of individual pT1SEstacker models and always achieved slightly better performance when voting cutoff was set as 0.6 (Fig. 6A; Supplementary Fig. S3).

**Fig 6.**
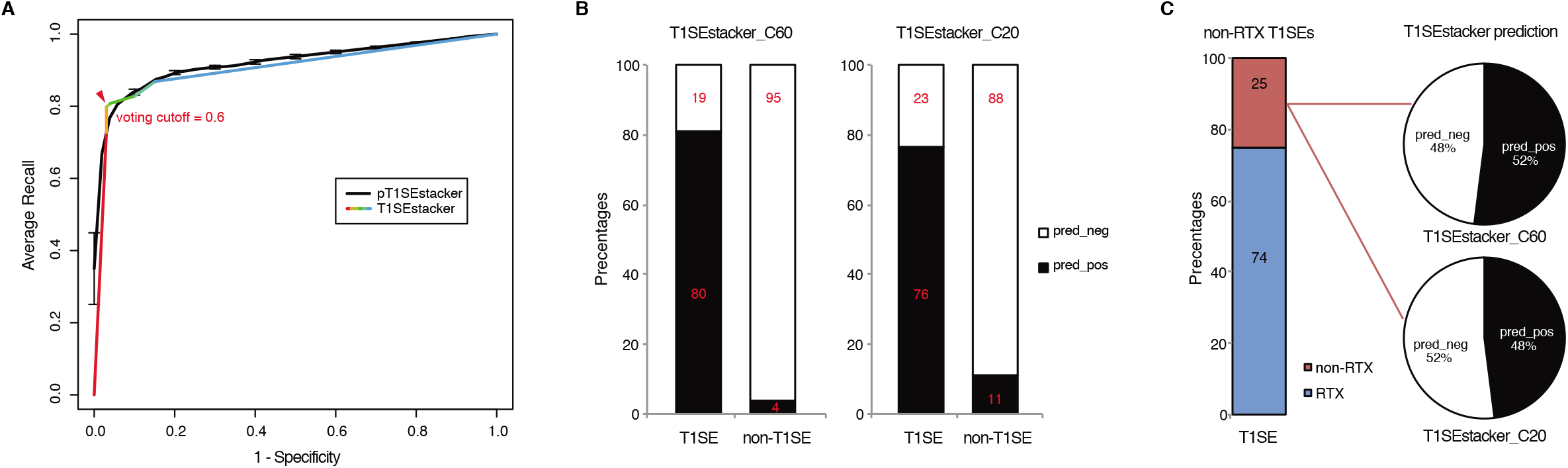
Prediction performance of T1SEstacker on experimentally verified T1SEs and the paired non-T1SEs. **(A)** ROC curves of the final T1SEstacker_C60 and the primary pT1SEstacker_C60 models on prediction of experimentally verified T1SEs and the paired non-T1SEs. The number for both T1SEs and non-T1SEs was 99. The best-optimized cutoff for the decision of T1SEstacker was indicated with a red arrow. **(B)** The recalling T1SEs and false positive T1SE predictions of T1SEstacker_C60 and T1SEstacker_C20. **(C)** The T1SE proteins without putative RTX motifs (non-RTX T1SEs) and the correctly predicted percentages of T1SEstacker on them.

### 8. T1SEstacker can recognize the common secretion signals among different types of T1SEs

We curated experimentally validated T1SEs, and applied the RTX protein prediction models to identify them. It should be noted that none of the C60 or C20 of the verified T1SEs contained any RTX motif. Both T1SEstacker_C20 and T1SEstacker_C60 could well predict the T1SEs (Fig. 6A; Supplementary Fig. S3). The recalling rate of T1SEstacker_C20 and T1SEstacker_C60 reached 77% and 81%, respectively (Fig. 6B). As a control, we used an independent negative dataset, and the specificity of T1SEstacker_C20 and T1SEstacker_C60 was 89% and 96%, respectively (Fig. 6B).

Among the validated T1SEs, 25% (25/99) do not contain any putative RTX motif along the full-length protein sequences (Fig. 6C; Supplementary Dataset S2). Interestingly, T1SEstacker_C60 correctly recalled 52% (13/25) of the non-RTX-motif T1SEs (Fig. 6C). Another one non-RTX-motif T1SE was also correctly recalled by at least pT1SEstacker_C60 models. The recalling rates of non-RTX-motif T1SEs are much higher than the false positive rates of the negative dataset for both C60 and C20 models (Fig. 6B-C). Therefore, the results further suggested that C-termini of T1SEs, with-RTX-motif or non-RTX-motif type, potentially contained common signals, which can guide the accurate prediction of these proteins.

### 9. Large variation of T1SE composition in *Salmonella* strains

The chromosomes of 26 representative strains from all *Salmonella* major tribes were scanned with T1SEstacker C60 model (Supplemental Dataset S3). In each strain, 269±22 T1SE candidates were predicted (Fig. 7A). With the recalling rate of 0.81 and false positive rate of 0.04 evaluated above on the validated T1SE dataset, the real number of T1SEs was estimated as 88 ~ 154, with an average of 123, in *Salmonella* strains (Fig. 7A). The precision of predicted T1SE candidates was only ~0.37 (123*0.81/269). However, it is difficult to improve the precision by shifting the decision cutoff values, or to distinguish the true positives from the false ones. Moreover, most of the real T1SEs were included in the predictions. Therefore, we used the original T1SEstacker predictions to analyze the distribution of T1SE candidates among the *Salmonella* strains.

**Fig 7.**
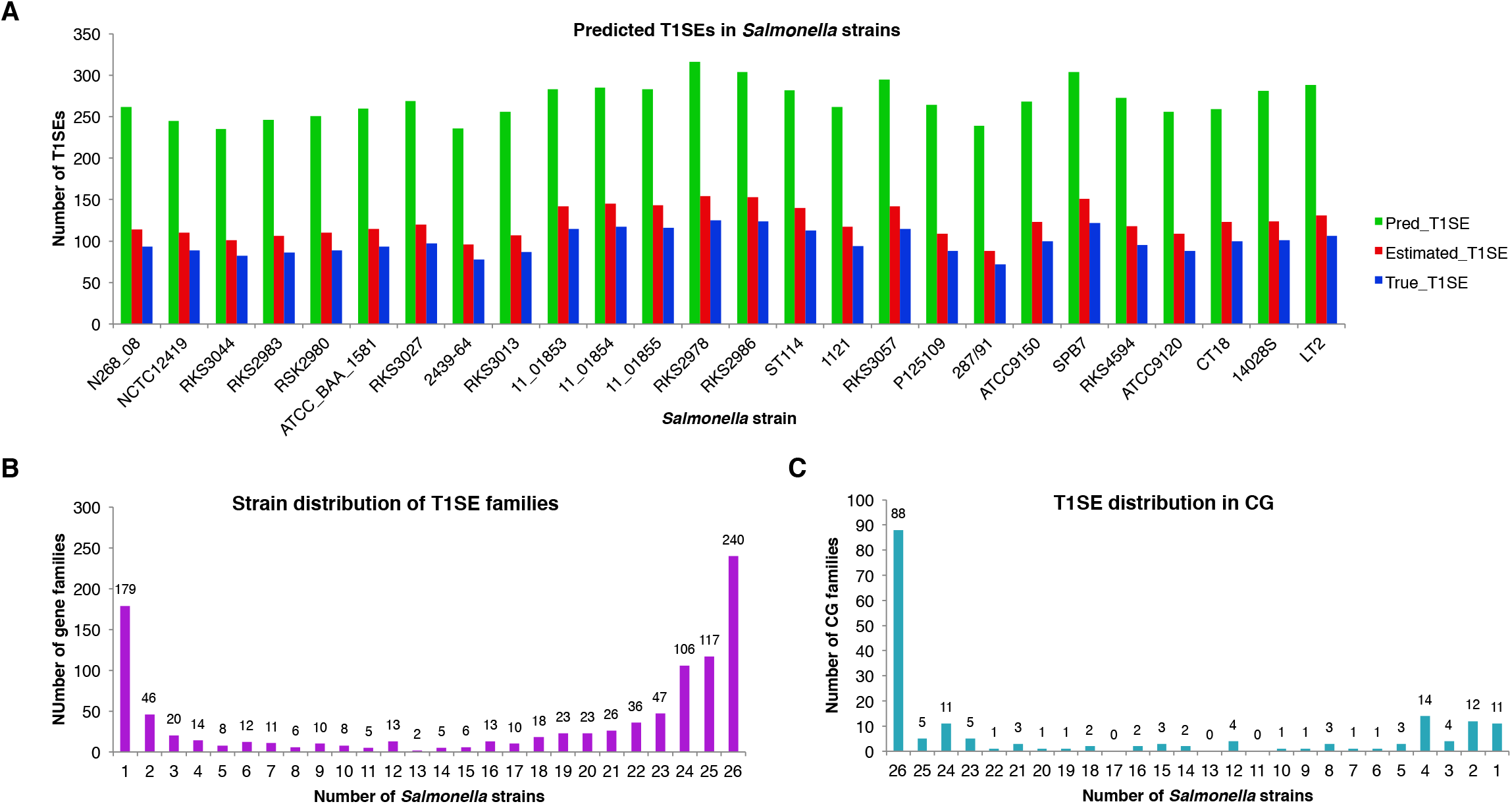
Distribution of T1SE candidates predicted from *Salmonella* strains with T1SEstacker. **(A)** Number of predicted T1SEs, estimated real T1SEs and true positive predictions in *Salmonella* strains. **(B)** The orthologous family distribution of T1SEs in *Salmonella* strains. **(C)** Distribution of core-genome T1SE hits in *Salmonella* strains. CG, core genome.

Despite a relatively stable number of T1SE candidates in different strains, the protein composition varied a lot. The candidates were clustered into 1,004 orthologous families, among which 240 (24%) were strain-specific proteins, 670 (67%) were present in fewer than half of the strains, and only 179 (18%) were distributed in the core genome of the *Salmonella* strains (Fig. 7B; Supplemental Dataset S3). For the core-genome hits, only 49% (88/179) were recognized as T1SEs in all the strains, and 31% (55/179) of the families were predicted as T1SEs only in fewer than a half of the strains (Fig. 7C; Supplemental Dataset S3). The results suggested that there is a large variety for the composition of T1SEs in different bacterial strains, and that a T1SE homolog does not necessarily remain a T1SE since mutations in the C-terminus could frequently avoid the recognition of T1SS.

## Discussion

Like other secreted proteins, bacterial type 1 secreted proteins (T1SEs) also play important roles in various infection diseases. Some T1SEs, e.g., bacteriocins, show non-self bacteria-killing activities and therefore have been used for anti-bacteria drug or probiotic development. How many T1SEs are there in each bacterial strain? How diverse is their function? The questions remain unanswered, since we are still at the very beginning on understanding the mechanisms of type 1 secretion. Only around 100 T1SEs have been verified by experiments, and many of them contain RTX motifs nearby the C-termini of protein sequences. However, not all T1SEs contain RTX motifs, while the proteins with RTX motifs, though more likely to be, are not necessarily T1SEs. Therefore, T1SEs could have other common targeted signals that mediate their specific type 1 secretion. More novel T1SEs could be identified based on these common signals.

Previous studies suggested possible signals within C-termini of RTX and non-RTX T1SEs. In this research, we focused on RTX T1SEs, observed the Aac features within their C-termini comprehensively, and compared them with the C-termini of non-RTX proteins or N-termini of the RTX and non-RTX proteins. It was interesting to identify specific Aac preference in C-termini of RTX proteins (Fig. 4A). As control, no apparent difference was found between the N-termini of RTX and non-RTX proteins (Fig. S1 and S2). The Aac preference profile was not biased by possibly included RTX motifs. On one hand, very few RTX motifs were retained in the observed length of C-terminal sequences (both C20 and C60) (Fig. 2C-D). On the other hand, the motif-enriched bi-AAs were not as strikingly different as other bi-AAs (Fig. 2B). Moreover, the real occurrence of some individual AAs or bi-AAs within C-termini of RTX proteins, especially C60, e.g., ‘G’ and ‘D’, was much higher than the percentage of proteins with putative RTX motifs within the region. Therefore, such Aac preference could be independent of RTX motif. Alternatively, RTX motifs could also represent the preference, but a more specific and conserved pattern. Besides the enriched Aac, significantly depleted Aac should also be noted, e.g., ‘E’, ‘K’, ‘R’ and ‘P’. In the research, by observing the position-specific Aac profiles, we also identified a typical amino acid composition pattern at the endmost C-termini of RTX proteins, with a motif feature of ‘[FLI][VAI]’ We also found that the C-termini of RTX proteins preferred β-strands rather than α-helices as in non-RTX proteins (Fig. 4B). It is intriguing to further investigate whether the unique amino acid composition and secondary structure contribute to the specificity of signal recognition of type 1 secretion.

Machine-learning models based on the C-terminal non-RTX-motif Aac features well-predicted RTX proteins from non-RTX proteins (Fig. 5; Table 2). The features within C20 showed certain power, while those buried in C60 showed better distinguishing capability (Fig. 5; Table 2). The C60 models could also accurately recall verified T1SEs at high prediction specificity (larger than 95%) (Fig. 6B). It should be pointed out again that, none of the verified T1SEs contained any RTX motif within C20 or C60 regions. More interestingly, 25 of the verified T1SEs do not contain RTX motif throughout their full-length sequences, and yet 12 and 13 were still predicted by C20 and C60 models, respectively, as positive results (Fig. 6C). Among the correctly predicted T1SEs, some are bacteriocins, and others are not putative RTX proteins. Therefore, the features identified in this study can be used for development of general T1SE prediction models. In future studies and as more non-RTX T1SEs have been identified, the common features can be reanalyzed, with a more balanced training dataset of different types of T1SEs.

We developed a tri-layer stacking model T1SEstacker, and showed that the stackers generally outperformed the individual machine-learning models (Table 2; Fig. 5C). Although some individual models also showed good performance, e.g., MM, RNN, SelfAttention and RF, but generally, not as good or stable as the stackers, pT1SEstacker (Table 2; Fig. 5C). We made a second round of stacking for the pT1SEstackers trained with sub-divided cross-validated datasets, because for pT1SEstackers we adopted a SVM model to integrating the prediction results of individual machine-learning models (Fig. 1). Similar with T1SEstacker that integrates pT1SEstacker results, other ensemblers often use voting strategy (Wang et al., 2019), or linearly weight each individual model (Hui et al., 2020). The parameters, i.e., linear weights for individual models and decision cutoffs for those models, were generally stable and not very sensitive to the sub-divided or full training datasets. However, for pT1SEstacker models, we trained the prediction results of individual models using SVM, and the parameters were pretty sensitive to the training datasets. Therefore, the 5 pT1SEstackers were each with different optimized parameters. To integrate their respective prediction results, another round of stacking had to be performed. The final model T1SEstacker appeared not apparently better than the pT1SEstacker models. However, once the optimized voting cutoff was selected (≥0.6, 3/5, consensus prediction), the prediction of T1SEstacker always showed best performance, with a compromise of sensitivity and specificity (Fig. 6A; Fig. S3).

The false positive rate (FPR) of T1SEstacker_C60 was low and close to 0.04. It is important, since many tools predicting bacterial secreted proteins showed a high FPR and the experimental research seldom benefited from the tool (Hui et al., 2020). As an example, we showed the influence of FPR on the final prediction performance, by prediction and estimation of T1SE candidates in *Salmonella* with T1SEstacker (Fig. 7A). Despite the high specificity (0.96), among the predicted T1SE candidates, a majority were false positives, and the precision was only ~0.37 (Fig. 7A). It is largely because for each genome most genes are non-T1SEs, and even 1% FPR could generate 50-100 false positive predictions, for which the number is close to that of true T1SEs. Therefore, it appears essential and urgent to further reduce FPR in predictor development, not merely for T1SE, but also for all types of secreted proteins.

Currently, there is still a lack of computational methods predicting T1SEs (Hui et al. 2021). Although Luo et al developed a random forest predictor, the tool or codes were not publically available and therefore a direct comparison could not be performed (Luo et al., 2015). An important factor that impedes development of prediction tools for T1SEs is the very limited number of experimentally validated T1SE proteins. Both Luo et al and we in this research used the Linhartova’s RTX proteins as the positive dataset (Luo et al., 2015; Linhartova et al., 2010). In fact, we also used the validated T1SEs to build a similar model, and the performance was only slightly inferior to T1SEstacker but the variance was much larger among the cross-validated replicates. Moreover, the T1SEstacker could accurately predict the novel ones in the validated effector dataset at a high specificity. Therefore, we presented the T1SEstacker based on Linhartova’s RTX proteins finally. With T1SEstacker and *Salmonella* strains, we also made estimation on the distribution of T1SEs. Roughly, there could be ~100 T1SEs in each bacterial strain. Therefore, the current T1SEs and function of T1SSs could be largely underestimated and underinvestigated. We also found the T1SE composition varied a lot among different bacterial strains, suggesting they could exert specific function for better fitting and bacterial survival. Therefore, it is of great significance to identify and investigate the function of T1SEs, for both microbiologists and computational biologists.

## Acknowledgement

The project was supported by the Funds for Medical Bioinformatics Youth Innovation Team of Shenzhen University (406/0000080805) and a Natural Science Fund of Shenzhen (JCYJ201607115221141). Z.C. and X.H. were supported by Special Funds for the Cultivation of Guangdong College Students’ Scientific and Technological Innovation, Climbing Program (pdjha0427). Z.C. was supported by a National Undergraduate Training Program of China for Innovation and Entrepreneurship (no. 201910590003).

## Supplementary Materials

Fig S1. Position-specific Aac profile difference between the C-termini of RTX proteins and three independent groups of non-RTX proteins.

Fig S2. Position-specific Aac profile difference between the N-termini of RTX proteins and three independent groups of non-RTX proteins.

**Fig S3. ROC curves of T1SEstacker and pT1SEstacker models on the verified T1SEs and non-T1SEs. (A)** Performance of T1SEstacker_C60 and pT1SEstacker_C60 on 99 verified T1SEs and 512 non-T1SEs. **(B)** Performance of T1SEstacker_C20 and pT1SEstacker_C20 on 99 verified T1SEs and paired 99 non-T1SEs. **(C)** Performance of T1SEstacker_C20 and pT1SEstacker_C20 on 99 verified T1SEs and 512 non-T1SEs. The best-optimized cutoff for the decision of T1SEstacker models were indicated with red arrows.

Dataset S1. Sequential Aac comparison between the C-termini of RTX and non-RTX proteins.

Dataset S2. RTX motif distribution within the experimentally verified T1SEs and the prediction results of T1SEstacker_C60 and T1SEstacker_C20.

Dataset S3. *Salmonella* T1SEs predicted with T1SEstacker.

